# Biological traits of seabirds predict extinction risk and vulnerability to anthropogenic threats

**DOI:** 10.1101/2020.09.30.321513

**Authors:** Cerren Richards, Robert S. C. Cooke, Amanda E. Bates

## Abstract

**Aim:** Here we aim to: 1) test whether globally-threatened vs non-threatened seabirds are separated in trait space; 2) quantify the redundancy and uniqueness of species trait combinations per IUCN Red List Category; and 3) identify traits that render species vulnerable to anthropogenic threats.

**Location:** Global

**Time period:** Contemporary

**Major taxa studied:** Seabirds

**Methods:** We compile and impute eight traits that relate to species’ vulnerabilities and ecosystem functioning across 341 seabird species. Using these traits, we build a mixed data PCA of species’ trait space. We further quantify trait redundancy with a unique trait combinations (UTCs) approach. Finally, we employ a similarity of percentages analysis (SIMPER) to identify which traits explain the greatest difference between threat groups.

**Results:** We find seabirds segregate in trait space based on threat status, indicating anthropogenic impacts are selectively removing large, long-lived, pelagic surface feeders with narrow habitat breadths. We further find that globally threatened species have higher trait redundancy, while non-threatened species have relatively unique ecological strategies and limited redundancy. Finally, we find that species with narrow habitat breadths, fast reproductive speeds, and omnivorous diets are more likely to be threatened by habitat-modifying processes (e.g., pollution and natural system modifications); whereas pelagic specialists with slow reproductive speeds and omnivorous diets are vulnerable to threats that directly impact survival and fecundity (e.g., invasive species and biological resource use).

**Main conclusions:** Our results suggest both globally threatened and non-threatened species contribute unique ecological strategies. Consequently, conserving both threat groups, but with contrasting approaches may avoid potential changes in ecosystem functioning and stability.

## INTRODUCTION

Humans are driving rapid changes in the world’s physical, chemical and biological makeup (Jenkins, 2003). Habitat transformation, species exploitation, climate change, pollution, and invasive species have the largest relative global impact (IPBES, 2019). These pressures are cumulative and have spread to all ecosystems, from the upper atmosphere to the deep sea (Bowler et al., 2020; Geldmann, Joppa, & Burgess, 2014; Halpern et al., 2008; Venter et al., 2016; Woolmer et al., 2008; Worm & Paine, 2016). Consequently, up to an estimated one million animal and plant species are now threatened with extinction (IPBES, 2019), populations of vulnerable taxa are declining, and biological diversity is changing (Dornelas et al., 2014).

Species traits are useful tools to understand species’ extinction risk, vulnerability to threats, and ecological roles (Peñaranda & Simonetti, 2015). Traits are attributes or characteristics of organisms measured at the individual level (Gallagher et al., 2020; Violle et al., 2007). Extinctions under human pressures are not random, but depend on a number of species’ traits such as body size, small geographic range, habitat specialisation, and slow life history (Cooke, Eigenbrod, & Bates, 2019; Davidson, Hamilton, Boyer, Brown, & Ceballos, 2009; Duffy, 2003; Gross & Cardinale, 2005; Peñaranda & Simonetti, 2015; Rao & Larsen, 2010). Therefore, threats likely impact ecologically similar species, while generalist traits, for example, large habitat breadths and generalist foraging strategies, may offer protection against extinction (Cooke, Eigenbrod, et al., 2019). Elucidating patterns and drivers of species’ extinction risk will likely provide the opportunity to develop more informed and effective conservation strategies (Ripple et al., 2017). Furthermore, selecting meaningful and interpretable species’ traits can relate to species’ vulnerability to threats and their contribution to ecosystem functions (Table 1). For example, a species’ diet captures regulation of trophic-dynamics and nutrient storage functions, and its sensitivity to changes at lower trophic levels (Tavares, Moura, Acevedo-Trejos, & Merico, 2019). Combinations of traits can summarise a species’ ecological role (Brum et al., 2017), and species can be grouped based on ecologically similar strategies (Cooke, Eigenbrod, et al., 2019).

**Table 1.** Description of the traits used in the present study and their relation to ecosystem functioning and species’ vulnerabilities. Ecosystem function column modified from Tavares et al. (2019). Imputation indicates the number of species imputed. Sources - 1: Cooke et al. (2019); 2: BirdLife International; 3: Wilman et al. (2014).

| Trait | Description | Imputed | Ecosystem Function | Species' Vulnerability | Source |
| --- | --- | --- | --- | --- | --- |
| Body Mass | Log<sub>10</sub> (median body mass in grams). | 0 | Nutrient storage and transport. | Strong predictor of extinction risk. | 1 |
| Habitat Breadth | Log<sub>10</sub> (number of IUCN habitats listed as suitable). | 0 | Nutrient transport. Community shaping through organism dispersal. | Exposure to threats across multiple locations, or limited one location. | 1 |
| Generation Length | Log<sub>10</sub> (generation length in years). | 1 | Nutrient storage. | The ability of populations to recover from threats. | 2 |
| Clutch Size | Number of eggs per clutch. | 4 | Nutrient storage. | The ability of populations to recover from threats. | 1 |
| Pelagic Specialism | Is the species a pelagic specialist?<br>*Pelagic Specialist*<br>*Non-pelagic Specialist* | 60 | Nutrient transport. | Exposure and interaction with marine threats, e.g. oil spills, bycatch. | 3 |
| Migration Strategy | Does migration occur?<br>*Full migrant*<br>*Non-migrant* | 3 | Nutrient transport. Community shaping through organism dispersal. | Exposure to threats across multiple locations, or limited one location. | 2 |
| Foraging Guild | The dominant foraging guild of the species.<br>*Diver*<br>*Surface*<br>*Ground*<br>*Generalist* | 60 | Nutrient storage. Trophic-dynamic regulations of populations. | The propensity of species to interact with threats, e.g. bycatch. | 3 |
| Diet Guild | The dominant diet of the species.<br>*Omnivore*<br>*Invertebrate*<br>*Vertebrate & scavenger* | 60 | Nutrient storage. Trophic-dynamic regulations of populations. | Sensitive to overexploitation of specific foods (e.g. overfishing) and changes in lower trophic levels. | 1 |

Seabirds are the most threatened group of birds and their status is deteriorating rapidly (Croxall et al., 2012; Paleczny, Hammill, Karpouzi, & Pauly, 2015). Seabirds are well adapted for life in the marine environment owing to their life history and ecological strategies including long life span, low fecundity and specialised foraging strategies e.g., diving for prey underwater. These traits likely evolved to optimise adult survival because delivering food to offspring from the open ocean requires large effort (Velarde, Anderson, & Ezcurra, 2019). However, seabirds require isolated terrestrial landmasses to breed during the breeding season. This requirement exposes seabirds to multiple and repeated anthropogenic threats in both the marine and terrestrial environment. These threats include those that directly affect survival and fecundity (e.g., invasive species, bycatch), threats that modify or destroy habitat (e.g., land modification, energy production) and global change threats (e.g., climate change) (Croxall et al., 2012; De Palma et al., 2015; Dias et al., 2019; Rodríguez et al., 2019).

Seabirds are an exceptionally well-studied faunal group and thus offer comprehensive biological detail for trait-based studies. Many seabird colonies are heavily monitored throughout the breeding season across the world. Furthermore, recent technological gains through miniaturization of biologging devices has revealed seabird behaviours at sea and during the winter (Richards, Padget, Guilford, & Bates, 2019). Thus, vast information is available on the life history, behavioural and ecological traits of seabirds. However, few studies have investigated the macroecological patterns of seabird threat risks. It remains an open question how ecological strategies of seabirds expose them to specific anthropogenic threats, and what consequence this has for ecosystem functioning.

Here we compiled and imputed eight traits across 341 seabird species to test whether species are separated in trait space based on extinction risk. We predict globally threatened species will occupy distinct regions of trait space because threats act on traits non-randomly (Duffy, 2003; Gross & Cardinale, 2005; Rao & Larsen, 2010). Next, we quantify the redundancy of species traits based on extinction risk (IUCN category). If pressures are targeting species with similar ecological strategies, we expect a greater redundancy in the traits of globally threatened species. Finally, we identify whether ecologically similar seabird species are responding similarly to human pressures. We hypothesize that species with narrow habitat breadths will be at risk from habitat modifying threats; species with slow reproductive speeds will be affected by pressures that directly affect survival and fecundity; and species with no threats will be generalists with fast reproductive speeds.

## METHODS

### Trait selection and data

We compiled data from multiple databases for eight traits (Table 1) across all 341 species of seabird. Here we recognise seabirds as those that feed at sea, either nearshore or offshore, but excluding marine ducks. These traits encompass the varying ecological and life history strategies of seabirds, and relate to ecosystem functioning and species’ vulnerabilities. We first extracted the trait data for body mass, clutch size, habitat breadth and diet guild from a recently compiled trait database for birds (Cooke, Bates, & Eigenbrod, 2019). Generation length and migration status were compiled from BirdLife International (datazone.birdlife.org), and pelagic specialism and foraging guild from Wilman et al. (2014). We further compiled clutch size information for 84 species through a literature search (see Appendix S1).

Foraging and diet guild describe the most dominant foraging strategy and diet of the species. Wilman et al. (2014) assigned species a score from 0 to 100% for each foraging and diet guild based on their relative usage of a given category. Using these scores, species were classified into four foraging guild categories (*diver*, *surface*, *ground*, and *generalist* foragers) and three diet guild categories (*omnivore, invertebrate*, and *vertebrate & scavenger* diets). Each was assigned to a guild based on the predominant foraging strategy or diet (score > 50%). Species with category scores ≤ 50% were classified as generalists for the foraging guild trait and omnivores for the diet guild trait. Body mass is the median body mass in grams. Habitat breadth is the number of habitats listed as suitable by the International Union for Conservation of Nature (IUCN, iucnredlist.org). Generation length describes the mean age at which a species produces offspring in years. Clutch size is the number of eggs per clutch. Migration status describes whether a species undertakes full migration or not. Pelagic specialism describes whether foraging is predominantly pelagic. To improve normality of the data, continuous traits, except clutch size, were log10 transformed.

### Multiple imputation

All traits had more than 80% coverage for our list of 341 seabird species, and body mass and habitat breadth had complete species coverage (Table 1). To achieve complete species trait coverage, we imputed missing data for clutch size (4 species), generation length (1 species), diet guild (60 species), foraging guild (60 species), pelagic specialism (60 species) and migration status (3 species). The imputation approach has the advantage of increasing the sample size and consequently the statistical power of any analysis whilst reducing bias and error (Kim, Blomberg, & Pandolfi, 2018; Penone et al., 2014; Taugourdeau, Villerd, Plantureux, Huguenin-Elie, & Amiaud, 2014).

We estimated missing values using random forest regression trees, a non-parametric imputation method, based on the ecological and phylogenetic relationships between species (Stekhoven & Bühlmann, 2012). This method has high predictive accuracy and the capacity to deal with complexity in relationships including non-linearities and interactions (Cutler et al., 2007). To perform the random forest multiple imputations, we used the *missForest* function from package “missForest” (Stekhoven & Bühlmann, 2012), based on 1,000 trees. We imputed missing values based on the ecological (the trait data) and phylogenetic (the first 10 phylogenetic eigenvectors, detailed below) relationships between species. Due to the predictive nature of the regression tree imputation approach, the estimated values will differ slightly each time. To capture this imputation uncertainty and to converge on a reliable result, we repeated the process 15 times, resulting in 15 trait datasets (González-Suárez, Zanchetta Ferreira, & Grilo, 2018; van Buuren & Groothuis-Oudshoorn, 2011). We take the mean trait values across the 15 datasets for subsequent analyses.

Phylogenetic information was summarised by eigenvectors extracted from a principal coordinate analysis, representing the variation in the phylogenetic distances among species (Jose Alexandre F. Diniz-Filho et al., 2012; José Alexandre Felizola Diniz-Filho, Rangel, Santos, & Bini, 2012). Bird phylogenetic distance data (Prum et al., 2015) were decomposed into a set of orthogonal phylogenetic eigenvectors using the *Phylo2DirectedGraph* and *PEM.build* functions from the “MPSEM” package (Guenard & Legendre, 2018). Here, we used the first 10 phylogenetic eigenvectors, ensuring a balance between including detailed phylogenetic information and diluting the information contained in the other traits. The first 10 eigenvectors in our data represented 61% of the variation in the phylogenetic distances among seabirds. Phylogenetic data can improve the estimation of missing trait values in the imputation process (Kim et al., 2018; Swenson, 2014). This is because closely related species tend to be more similar to each other (Pagel, 1999) and many traits display high degrees of phylogenetic signal (Blomberg, Garland, & Ives, 2003). While imputation error is minimised when including the first 10 phylogenetic eigenvectors as variables in the imputations (Penone et al., 2014), these phylogenetic eigenvectors are more representative of divergences closer to the root of the phylogeny and do not include fine-scale differences among species (Jose Alexandre F. Diniz-Filho et al., 2012).

To quantify the average error in random forest predictions across imputed datasets (out-of-bag error), we calculated the normalized root mean squared error for continuous traits (clutch size = 13%, generation length = 0.6%) and percent falsely classified for categorical traits (diet guild = 29%, foraging guild = 18%, pelagic specialism = 11%, migration status = 19%). Since body mass and habitat breadth have complete trait coverage, their out-of-bag error is 0%. Low imputation accuracy is reflected in high out-of-bag error values where diet guild had the lowest imputation accuracy with 29% wrongly classified on average.

### Sensitivity

To compare whether our results and conclusions were quantitatively and qualitatively similar between the imputed and non-imputed datasets, we ran all of our analyses with and without the imputed data.

### Species extinction risk

The International Union for Conservation of Nature’s (IUCN) Red List of Threatened Species (iucnredlist.org) is the most comprehensive information source on the global conservation status of biodiversity (IUCN, 2020). This powerful tool classifies species into nine categories of extinction risk. Here we use five IUCN Red List categories to group extant species into broader global risk groups. Species categorised as Critically Endangered (CR), Endangered (EN) and Vulnerable (VU) were defined as *globally threatened*, and species classified as Near Threatened (NT) and Least Concern (LC) were defined as *non-threatened*.

Two species classified as Data Deficient (*Oceanites gracilis* and *Oceanitespincoyae*) and one Not Evaluated species (*Larus thayeri*) were removed from the species list leaving a total of 338 species for all subsequent analyses.

### Principal component analysis of mixed data

To quantify the trait space shared by globally threatened and non-threatened seabirds, we ordinated 338 seabirds based on eight traits with a principle component analysis (PCA) of mixed data. We used the package “PCAmixdata” and function *PCAmix* (Chavent, Kuentz, Labenne, Liquet, & Saracco, 2017). PCA of mixed data takes a two-step approach through merging the standard PCA with multiple correspondence analysis (MCA) (Chavent, Kuentz-Simonet, Labenne, & Saracco, 2014). For continuous data, PCAmix is a standard PCA, whereas for categorical data, PCAmix it is an MCA (Chavent et al., 2014). To quantify the degree to which threat status explains trait space variations among seabirds, we used the permutational MANOVA framework in the *adonis* function and package “vegan” (Oksanen et al., 2018).

### Trait-level distributions and proportions

To test whether the traits of globally threatened and non-threatened seabirds are different at the individual trait level, we explored the distributions of continuous traits and proportions of categorical traits per threat category. Differences in the means of threatened and non-threatened species within continuous traits were compared with Mann-Whitney U tests using function *wilcox.test* (R Core Team, 2018). We further calculated Hedge’s g effect size with function *hedges_g* and package ‘effectsize’ (Ben-Shachar, Makowski, & Lüdecke, 2020). For categorical traits, we tested for independence with a Chi-squared approach using function *chisq.test* (R Core Team, 2018).

### Unique trait combinations

To quantify the redundancy and uniqueness of species trait combinations per IUCN Red List Category, we used unique trait combinations (UTCs). Here UTC is defined as the proportion of species with trait combinations that are not found in other seabird species. To compute the UTCs of the 338 seabirds, we broke the continuous traits into three equally spaced bins (small, medium and large) between minimum and maximum values. Following this, the proportion of UTCs within each IUCN Red List Category was calculated as a percentage.

### Seabird Threats

We extracted the past, present, and future threats for 338 seabirds from the IUCN Red List database using the function *rl_threats* and package “rredlist” (Chamberlain, 2018). These data have recently been updated in a quantitative review from >900 publications (Dias et al., 2019), and are classified into 12 broad types (Table 2). We reclassified the IUCN threats into four general categories (Table 2): (1) *direct* – threats that directly affect survival and fecundity; (2) *habitat* - threats that modify or destroy habitat; (3) *no threats –* species with no identified IUCN threats; and (4) *other* – threats that are indirectly or not caused by humans (Gonzalez-Suarez, Gomez, & Revilla, 2013). We excluded *other* threats (climate change and severe weather, and geological events) from our analyses because they are not directly linked to anthropogenic activity.

**Table 2.**
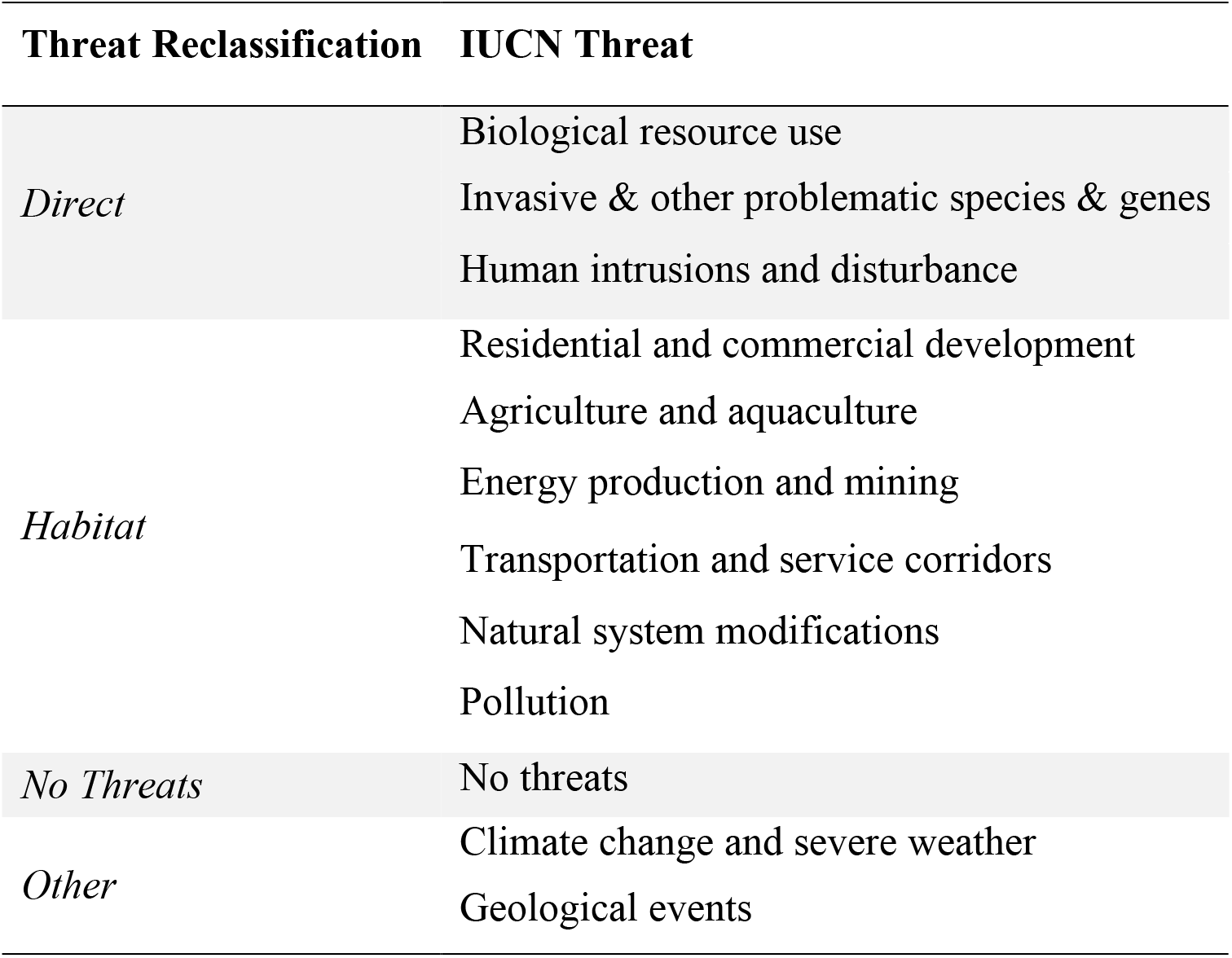
IUCN reclassified threat categories. ‘Direct’ threats directly affect survival and fecundity. ‘Habitat’ threats modify or destroy habitat. ‘No threats’ encompasses species with no identified IUCN threats. ‘Other’ threats are indirectly or not caused by humans. Modified from Gonzalez-Suarez, Gomez & Revilla (2013).

### SIMPER analysis

To identify which traits explain the greatest difference between threats, we took a similarity of percentages (SIMPER) approach using the function *simper* in package “vegan” (Oksanen et al., 2018). SIMPER typically identifies the species that contribute the greatest dissimilarity between groups (levels) by disaggregating the Bray-Curtis similarities between inter-group samples from a species-abundance matrix (Clarke & Warwick, 2001). Here, we assembled a trait-by-threat matrix, where traits are each level of the categorical and binned continuous traits (23 levels) and threats are the IUCN threat categories (first 10 levels from Table 2). For each threat, we calculated the proportion of species in each trait category. The reclassified IUCN threats were used to isolate the traits that contribute the greatest difference between habitat threats, direct threats and no threats.

All analyses were performed in R version 3.5.0 (R Core Team, 2018).

## RESULTS

### Threat status segregation in multidimensional trait space

We find globally threatened species are distinct from non-threatened species in terms of their biological trait diversity (PERMANOVA, R^2^ = 0.122, p = 0.001; Fig. 1). Together, the first two dimensions (identified herein as “Dim1” and “Dim2”) of the mixed data PCA explain 41% of the total trait variation (Fig. 1). Dim1 integrates reproductive speed, the trade-off between clutch size (loading = 0.860) and generation length (loading = −0.696), invertebrate diet (loading = 0.645), vertebrate & scavenger diet (loading = −0.881), omnivore diet (loading = −0.158), pelagic specialism (loading = −0.306), non-pelagic specialism (loading = 1.336) and surface foragers (loading = −0.855). Species with high Dim1 scores are typically characterised as non-pelagic scavengers with fast reproductive speeds e.g., cormorants, gulls and terns. Species with low Dim1 values have slow reproductive speeds and are pelagic surface foragers with diets high in invertebrates e.g., albatross, petrels, shearwaters and storm-petrels. Dim2 integrates body mass (loading = −0.347), full migrants (loading = 0.360), non-migrants (loading = −1.088), divers (loading = −0.967), generalists (loading = 0.979) and ground (loading = 1.481) foraging strategies. Species with high Dim2 are small bodied ground or generalist foragers e.g., gulls, terns, skuas and jaegers while those with low Dim2 are large bodied non-migrating divers e.g., shags, boobies and penguins.

**Figure 1.**
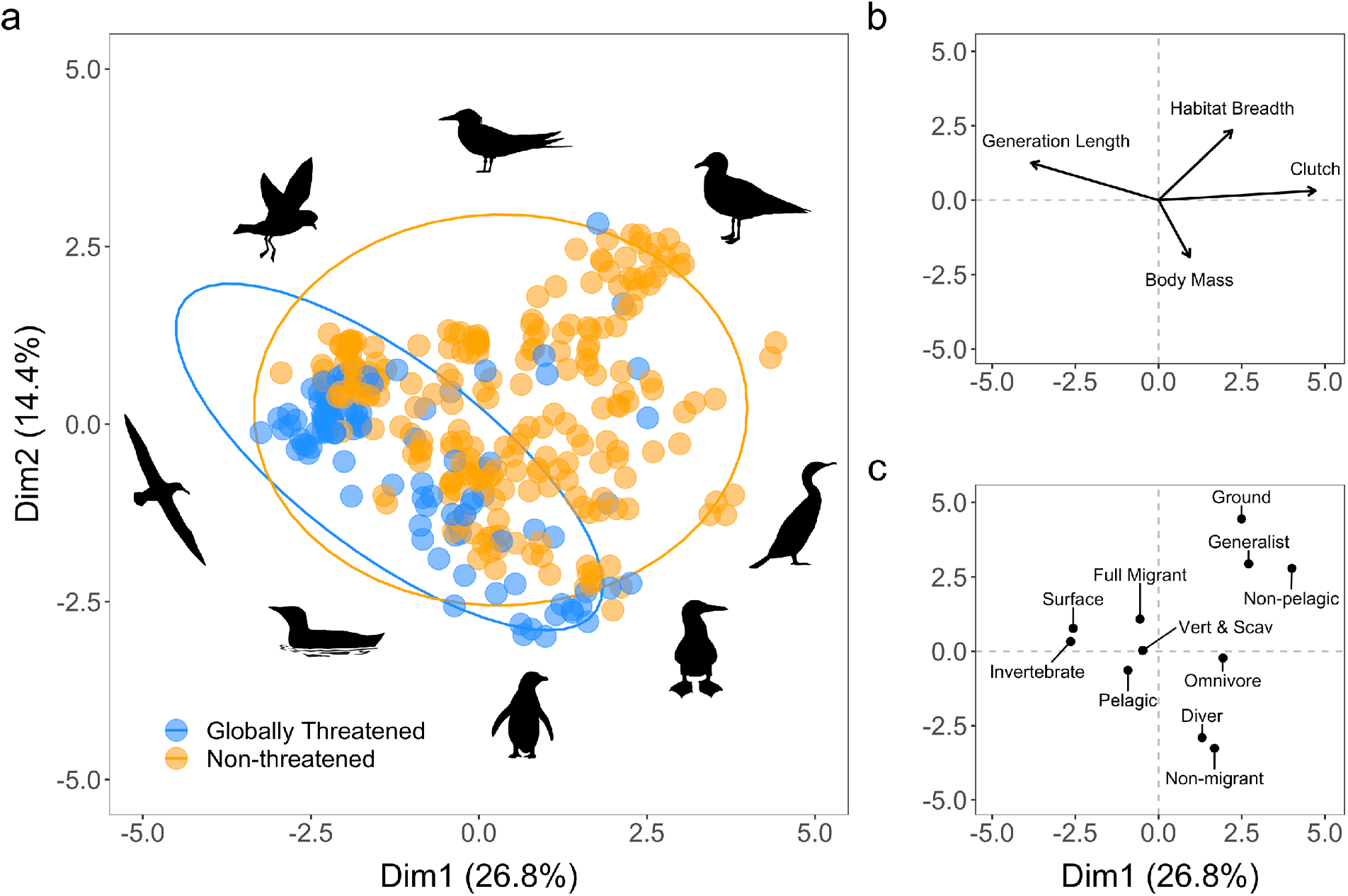
Mixed data PCA biplot of seabird traits. a) Points are the principal component scores of each seabird (mean values across 15 imputed datasets). Ellipses indicate the 95% confidence intervals for globally threatened (blue) and non-threatened (orange) seabird species. Silhouettes represent a selection of families aggregated at the edge of trait space. All silhouettes created by authors. Coordinates of b) continuous and c) categorical traits. Coordinates were rescaled to match the mixed data PCA.

Ten species fall outside the 95% confidence interval ellipse for globally threatened species. These include eight Laridae (Black-billed Gull, Black-fronted Tern, Relict Gull, Black-bellied Tern, Chinese Crested Tern, Indian Skimmer, Aleutian Tern, Lava Gull), one Phalacrocoracidae (Chatham Islands Shag) and one Spheniscidae (Galapagos Penguin).

### Individual trait differences

We find a significant difference in six traits between globally threatened and non-threatened species (Fig. 2; Table 3). Specifically, habitat breadths of globally threatened species are 2.2 times smaller [95% CI: −2.52, −1.95] than non-threatened seabirds, clutch sizes are 0.46 times smaller [95% CI: −0.69, −0.22], and generation lengths are 0.43 times longer [95% CI: 0.20, 0.67]. Compared to non-threatened species, we find globally threatened species have 18.8% more pelagic specialists, 26.5% more surface foragers, 5.0% fewer divers, 4.2% fewer ground foragers, 17.4% fewer generalist foragers, 31.5% fewer species with invertebrate diets, 22.5% greater species with vertebrate and scavenger diets, and 9.0% greater species with omnivore diets (Fig 2). There was no difference in the body mass, 0.24 times larger [95% CI: 0.01, 0.47], or migration traits (0.13% greater full migrants) between globally and non-threatened species (Table 3). We therefore find globally-threatened species are typically surface feeders with a diet higher in fish and carrion. They are mostly pelagic specialists that have narrow habitat breadths, small clutch sizes and long generation times. In comparison, non-threatened species are typically generalist foragers with a diet high in invertebrates. These species also typically have shorter generation lengths and larger clutch sizes with a broader habitat breadth and less pelagic specialism.

**Figure 2.**
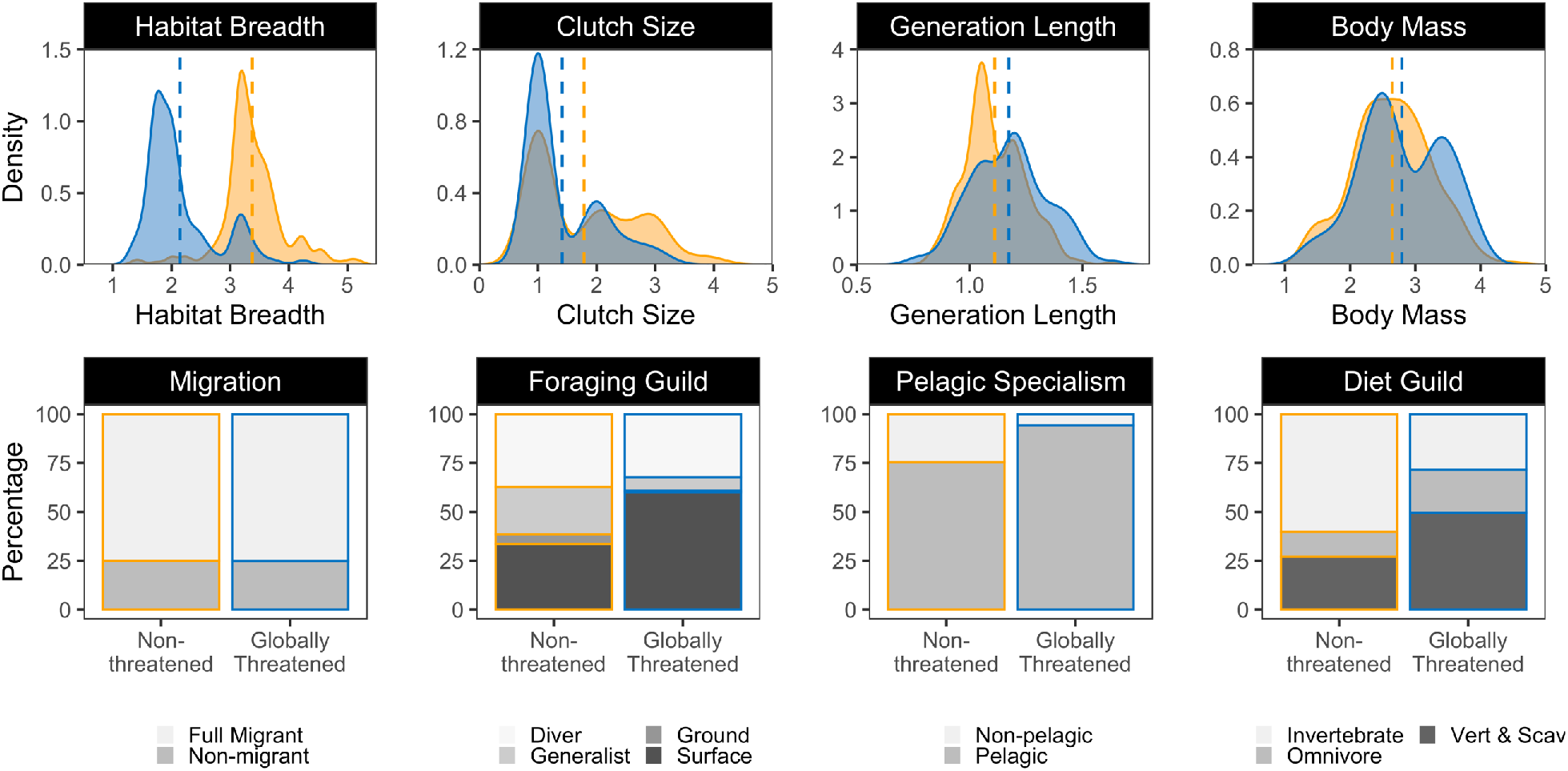
Distributions of continuous traits and proportion of categorical traits. Orange represents non-threatened species, while blue represents globally threatened species. Dashed lines are the mean of each distribution. Habitat breadth, generation length and body mass x-axes are log-transformed.

**Table 3.** Output results from the Mann-Whitney U and Chi-Squared tests which test the difference in the means (Mann-Whitney U) and independence (Chi-Squared) between the traits of globally threatened and non-threatened species.

| **Continuous Trait** | **Mann-Whitney U (W)** | **p-value** |
| --- | --- | --- |
| Body Mass | 13814 | 0.0571 |
| Clutch Size | 9431 | 0.0002 |
| Habitat Breadth | 2077.5 | 0.0000 |
| Generation Length | 15187 | 0.0003 |
| **Categorical Trait** | **Chi-squared (X<sup>2</sup>)** | **p-value** |
| Diet Guild | 28.812 | 0.0000 |
| Pelagic Specialism | 15.565 | 0.0000 |
| Foraging Guild | 27.733 | 0.0000 |
| Migration Strategy | 1.4119e<sup>-30</sup> | 1.0000 |

### Trait redundancy and uniqueness

We classify 166 different trait combinations across 338 seabirds. Of these, 59% are composed of only one species (n = 98) and are defined as unique trait combinations (UTCs). The proportion of UTCs decreases with increasing IUCN threat level (Fig 3). Consequently, a greater proportion of non-threatened species (32%) contribute UTCs than globally threatened species (23%). We, therefore, find greater redundancy in traits of globally threatened species and greater uniqueness in traits of non-threatened species (Fig. 3).

**Figure 3.**
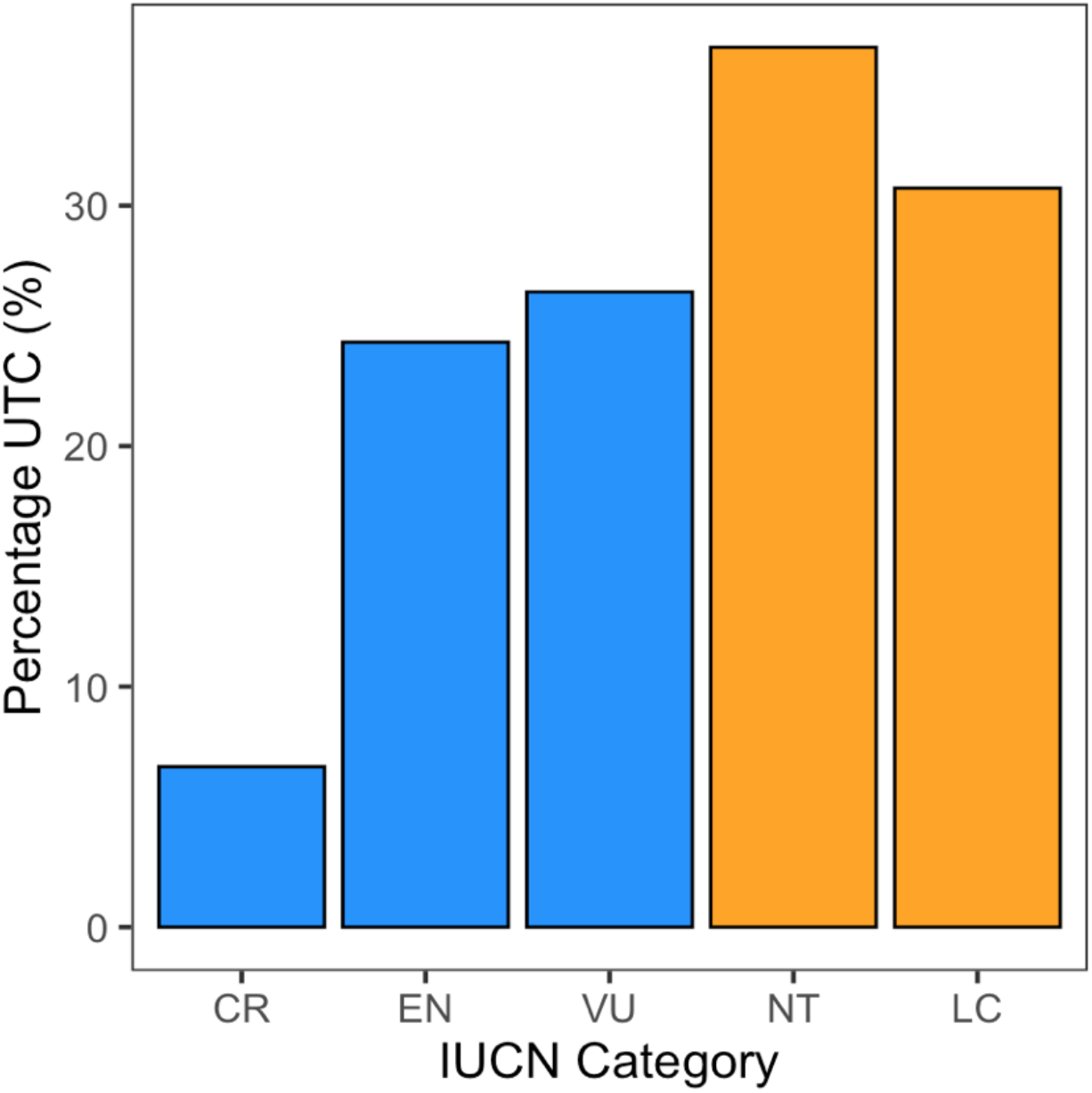
Proportion of seabird species with unique trait combinations (UTC) for each IUCN category. Orange represents non-threatened categories and blue represents globally threatened categories. ‘CR’ is Critically Endangered, ‘EN’ is Endangered, ‘VU’ is Vulnerable, ‘NT’ is Near Threatened, and ‘LC’ is Least Concern.

### SIMPER

Similarity percentages analysis (SIMPER) identifies the combination of reproductive speed traits (generation length and clutch size) and specialisation traits (pelagic specialism, diet guild, and habitat breadth) drive the greatest dissimilarity between threat types (Table 4). Specifically, we find that reproductive speed and pelagic specialism traits drive the greatest dissimilarity between direct and habitat threats. Diet, pelagic specialism, and habitat breadth traits explain the greatest dissimilarity between direct threats and no threats. Finally, diet, habitat breadth, and generation length traits explain the greatest dissimilarity between habitat threats and no threats.

**Table 4.** SIMPER summary of top five traits contributing to the Bray Curtis dissimilarity between threats. The proportion of species per trait is indicated as greater (+), or smaller (-) between each threat category. ‘S’ indicates ‘small’ and ‘M’ indicates ‘medium’.

| Threat Contrast | Trait | Contribution (%) | Cumulative (%) | Direct | Habitat | No Threat |
| --- | --- | --- | --- | --- | --- | --- |
| *Direct vs. Habitat* | Non-pelagic Specialism | 7.3 | 7.3 | - | + |  |
|  | Pelagic Specialism | 7.3 | 14.7 | + | - |  |
|  | Clutch Size (S) | 7.3 | 22.0 | + | - |  |
|  | Generation Length (S) | 7.0 | 29.0 | - | + |  |
|  | Generation Length (M) | 6.0 | 35.0 | - | + |  |
| *Direct vs. No Threats* | Omnivore Diet | 9.8 | 9.8 | + |  | - |
|  | Non-pelagic Specialism | 7.5 | 17.3 | - |  | + |
|  | Pelagic Specialism | 7.5 | 24.8 | + |  | - |
|  | Invertebrate Diet | 7.3 | 32.0 | - |  | + |
|  | Habitat Breadth (S) | 7.2 | 39.2 | + |  | - |
| *Habitat vs. No Threats* | Omnivore Diet | 8.6 | 8.6 |  | + | - |
|  | Habitat Breadth (S) | 8.1 | 16.6 |  | + | - |
|  | Habitat Breadth (M) | 7.9 | 24.5 |  | - | + |
|  | Invertebrate Diet | 7.6 | 32.1 |  | - | + |
|  | Generation Length (S) | 6.2 | 38.3 |  | - | + |

Through generalising the directionality of important trait contributors between each threat (Table 4), we find seabirds with different traits are at risk from direct and habitat threats (Fig. 4). Species with slow reproductive speeds, specialisation traits (pelagic specialism) and omnivorous diets are at greater risk from direct threats. Direct threats target all families of seabird, but most are tubenose seabirds (albatross, shearwaters, and petrels). Habitat threats typically endanger those with fast reproductive speeds, specialisation traits (small habitat breadth) and omnivorous diets. These species are typically gulls and terns, yet habitat threats target all families of seabird. Species with no threats, which are primarily gulls, have the fastest reproductive speeds, generalist traits (non-pelagic specialism, larger habitat breadth) and invertebrate diets.

**Figure 4.**
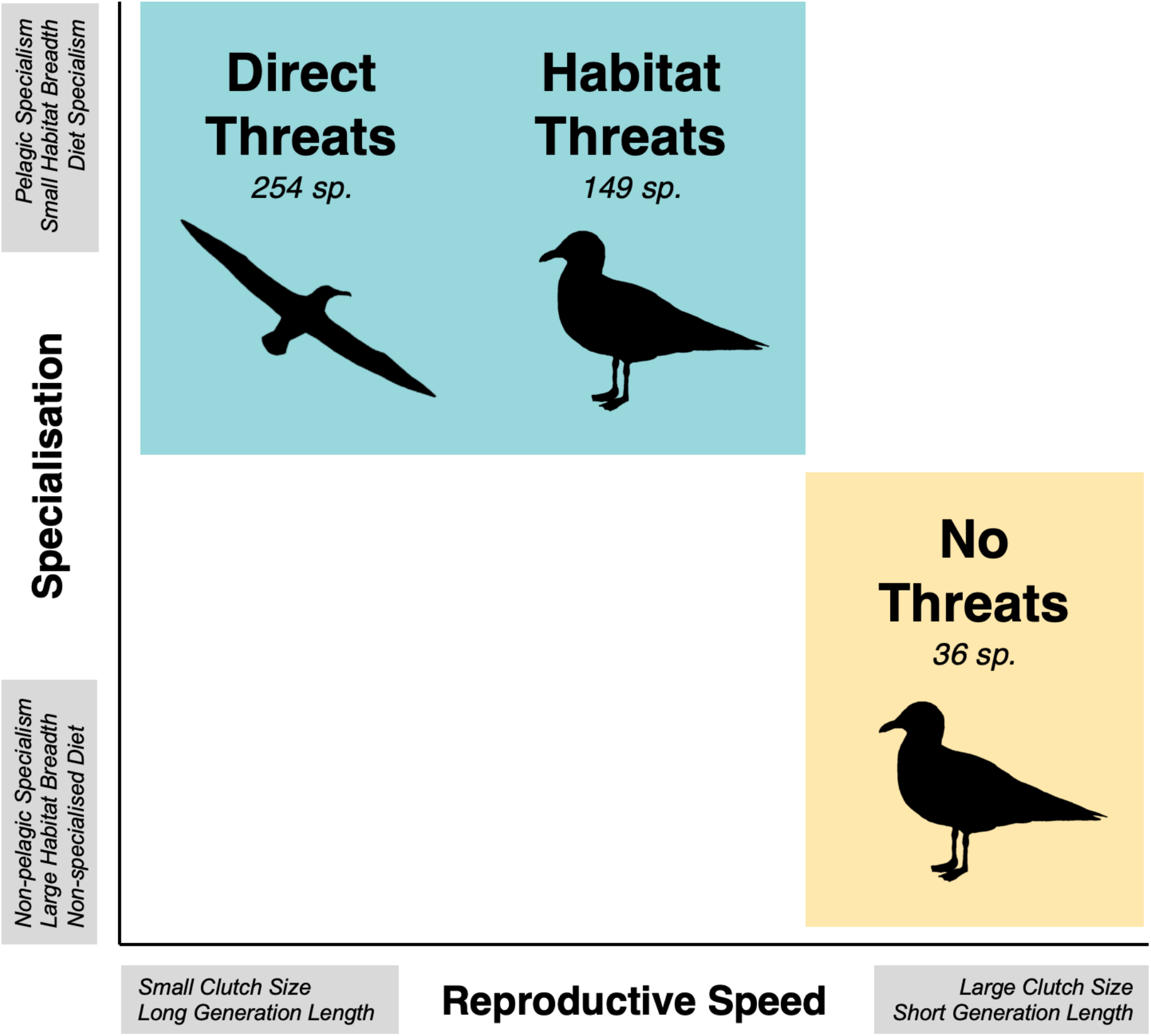
Generalised pattern of traits that predict vulnerability of seabirds to varying anthropogenic threats based on the results presented in Table 4. Silhouettes represent seabird families with high frequencies of species at risk to each threat type. ‘Direct’ threats directly impact the survival and fecundity of seabirds, while ‘habitat’ threats modify or destroy habitats. ‘No threats’ encompasses species with no identified IUCN threats. Reproductive speed is the trade-off between clutch size and generation length. Specialisation encompasses pelagic specialism, habitat breadth and diet guild.

### Sensitivity

We find that our results and conclusions are comparable between the imputed and non-imputed datasets (see Appendix S2 in supporting information).

## DISCUSSION

We reveal that globally threatened and non-threatened seabirds occupy different regions of trait space. Specifically, globally threatened species share a distinct subset of similar traits that are associated with a higher risk of extinction. Therefore, the loss of threatened species, such as wide-ranging albatross and shearwaters, may have direct implications for ecosystem functioning such as trophic regulation, nutrient transportation and community shaping (Graham et al., 2018; Tavares et al., 2019). We further find non-threatened species have relatively unique ecological strategies and limited redundancy. Consequently, non-threatened species may have less insurance to buffer against ecosystem functioning declines should they become threatened in the future (Yachi & Loreau, 1999). We must therefore prioritise the conservation of both globally threatened and non-threatened species, but with contrasting approaches to avoid potential changes in ecosystem functioning and stability. Globally threatened species would benefit from targeted conservation interventions, whereas non-threatened species require long-term monitoring of their populations and environment (e.g. Hebert et al., 2020).

We find a number of traits emerge with strong association to extinction risk and different types of threats. Overall, anthropogenic pressures may be selecting against slow-lived and specialised species, e.g., albatross and petrels, in favour of fast-lived and wide-ranging generalists, e.g., gulls and terns. This agrees with the patterns of other birds and mammals (Cooke, Eigenbrod, et al., 2019; Davidson et al., 2009; Peñaranda & Simonetti, 2015). However, in contrast to numerous studies (Cardillo et al., 2005; Cooke, Eigenbrod, et al., 2019; Ripple et al., 2017), we find no difference in the body mass of globally and non-threatened species. Therefore, threats are indiscriminate across seabirds from the largest (Wandering Albatross, *Diomedea exulans*, 7000 g) to the smallest seabird (European Storm-petrel, *Hydrobates pelagicus*, 25 g). Potential explanations could be that major threats to seabirds are not size dependent. For example, invasive species on a breeding island would consume all species’ eggs, and all sizes of seabirds are attracted to fishing vessels. Moreover, large seabirds are less targeted for hunting in comparison to mammals, e.g., game mammals.

Traits distinguishing species at risk from direct threats were slow reproductive speed, pelagic specialism, and diet guild traits, reflecting recent findings for mammals (Gonzalez-Suarez et al., 2013). Here, direct threats encompass invasive species and bycatch, which are the top two threats facing seabirds worldwide (Dias et al., 2019), in addition to human disturbance. Most species at risk to direct threats are tubenose seabirds (albatross, petrels, shearwaters). Tubenoses are highly pelagic species that depend on the ocean for foraging. Therefore, tubenoses often strongly overlap with fishing vessels (Clay et al., 2019) and opportunistically scavenge fisheries discards. In this process, birds are caught on baited hooks and drowned, or entanglement in nets and collide with cables which results in high mortality. Consequently, an estimated 320,000 seabirds die annually in longline fleets alone (Anderson et al., 2011). Tubenose seabirds are further strongly impacted by invasive species (e.g., rats and cats) and human disturbance at breeding colonies. These seabirds lay a single egg per season; therefore, their populations have a lower capacity to compensate for bycatch mortality and poor reproductive success due to invasive species and human disturbance.

We find species at risk to habitat threats have the smallest habitat breadths, and slower reproductive speeds than species with no threats, and omnivorous diets. This finding corroborates previous studies which identify habitat specialisation increases species’ vulnerability and limits their capacity to adapt to environmental change (Gonzalez-Suarez et al., 2013; Peñaranda & Simonetti, 2015). Habitat threats particularly target species such as cormorants and gulls. This is likely because coastal and wetland habitats are vital for these seabirds during wintering and breeding, yet they are being modified and destroyed by activities such as land use change and tourism.

Identifying traits most associated with threats can lead to more informed and effective conservation strategies. Species at risk to direct threats need targeted conservation interventions through bycatch mitigation and invasive species eradication to protect highly pelagic species with slow reproductive speeds. These initiatives are beginning to show great promise. For example, implementing bird deterrents in a South African trawl fishery reduced albatross deaths by 95% between 2004 to 2010 (Maree, Wanless, Fairweather, Sullivan, & Yates, 2014). Furthermore, eradicating rats from breeding colonies has dramatically recovered seabird populations (Veitch et al., 2019), and restored ecosystem functions such as nutrient transportation to soil and plants (Jones, 2010; Wardle, Bellingham, Bonner, & Mulder, 2009; Wardle, Bellingham, Fukami, & Bonner, 2012). Habitat breadth is strongly related to threat status, therefore many species will benefit from habitat conservation. For example, through designating protected areas at sea to conserve important seabird hotspots, movement pathways and foraging areas (D’Aloia et al., 2019; Ronconi, Lascelles, Langham, Reid, & Oro, 2012). At breeding sites, closing colony visitation during the breeding season and establishing buffer zones for land, water, and air could eliminate disturbance and nest abandonment.

Here we use the IUCN database to identify the traits most associated with different threats. While the IUCN threats is a valuable resource, its collation via expert opinion is subjective and can contain bias (Hayward, 2009), therefore threats may be unreported or overreported. Furthermore, rare or understudied species, for example the Critically Endangered magenta petrel (*Pterodroma magenta*) with fewer than 100 mature individuals, likely have fewer known threats than highly studied species such as the Atlantic puffin (*Fratercula arctica*). Further studies that couple spatial patterns of extrinsic threats with intrinsic traits could offer valuable insight into species vulnerabilities to anthropogenic threats, and ultimately help inform effective management and conservation at local and global scales.

In conclusion, we expand our understanding of extinction risk drivers in seabirds through a trait-based approach. Our findings highlight the need to conserve both globally and non-threatened species in order to conserve the diversity of ecological strategies and associated ecosystem functions. We suggest traits be coupled with spatial patterns of extrinsic threats to advance conservation management strategies.

## Data Accessibility Statement

The raw (non-imputed) trait data are provided as Appendix S3. The complete (mean imputed) trait data used for all analyses are provided as Appendix S4.

## Acknowledgements

We thank Dave Fifield and Shawn Leroux for comments and suggestions that greatly improved the manuscript.

