## Appendix S2 for "Biological traits of seabirds predict extinction risk and vulnerability to anthropogenic threats"

### Appendix S2: Sensitivity Analysis

To compare whether our results and conclusions were quantitatively and qualitatively similar between the imputed and non-imputed datasets, we ran all of our analyses with and without the imputed data.

#### Mixed data PCA

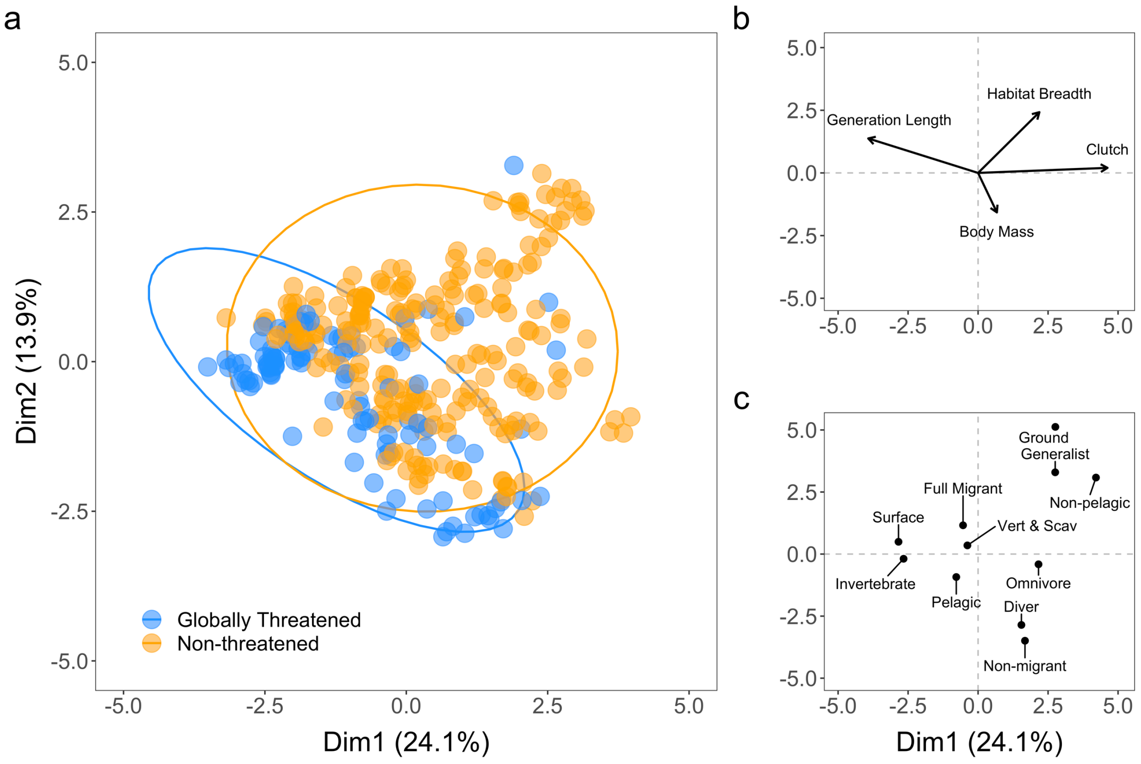


*Figure S2.1 Mixed data PCA biplot of seabird traits excluding imputed data. a) Points are the principal component scores of each seabird species. Ellipses indicate the 95% confidence intervals for globally threatened (blue) and non-threatened (orange) seabird species. Coordinates of b) continuous and c) categorical traits. Coordinates were rescaled to match the mixed data PCA.*

```
Call:
adonis(formula = pca_permanova ~ traits$Threat, method = "euclidean")

Permutation: free
Number of permutations: 999

Terms added sequentially (first to last)

               Df SumsOfSqs MeanSqs F.Model      R2 Pr(>F)    
traits$Threat   1    182.25 182.253  47.148 0.12305  0.001 ***
Residuals     336   1298.82   3.866         0.87695           
Total         337   1481.07                 1.00000           
---
Signif. codes:  0 ‘***’ 0.001 ‘**’ 0.01 ‘*’ 0.05 ‘.’ 0.1 ‘ ’ 1
```

*Figure S2.2 R output that quantifies the degree to which threat status explains trait space variations among seabirds.*

### Individual Trait differences

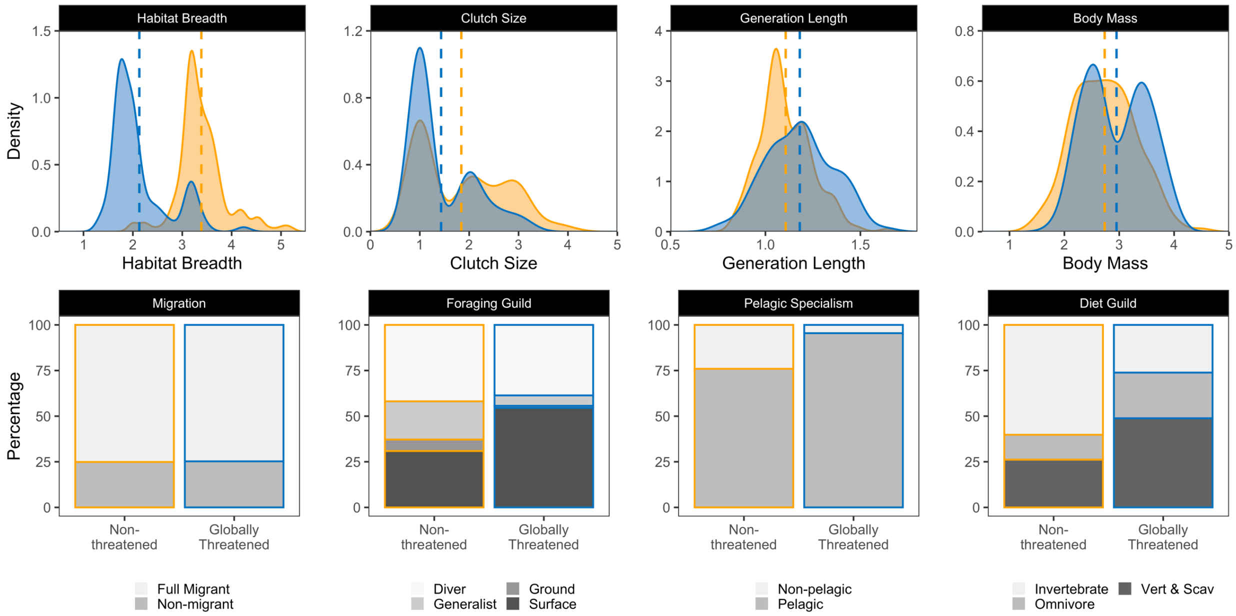


*Figure S2.3 Distributions of continuous traits and proportion of categorical traits, excluding imputed data. Orange represents non-threatened species, while blue represents globally threatened species. Habitat breadth, generation length and body mass x-axes are log-transformed. Habitat breadth and body mass have full trait coverage.*

*Table S2.1 Output results from the Mann-Whitney U and Chi-Squared tests which test the difference in the means (Mann-Whitney U) and independence (Chi-Squared) between the non-imputed traits of threatened and non-threatened species. Habitat breadth and body mass have full trait coverage.*

| **Continuous Trait** | **Mann-Whitney U (W)** | **p-value** |
|---|---|---|
| Body Mass | 13814 | 0.0571 |
| Clutch Size | 8962 | 0.0001 |
| Habitat Breadth | 2077.5 | 0.0000 |
| Generation Length | 15003 | 0.0004 |
| **Categorical Trait** | **Chi-squared ($X^2$)** | **p-value** |
| Diet Guild | 27.982 | 0.0000 |
| Pelagic Specialism | 14.335 | 0.0000 |
| Foraging Guild | 21.069 | 0.0001 |
| Migration Strategy | 0 | 1.0000 |

*Table S2.2 Hedges g effect size and 95% confidence intervals for non-imputed continuous traits, and the percentage difference between non-imputed categorical traits of globally threatened and non-threatened species. Habitat breadth and body mass have full trait coverage.*

| **Continuous Trait** | **Effect Size** | **95% CI** |
|---|---|---|
| Body Mass | 0.24 | [0.01, 0.47] |
| Clutch Size | -0.48 | [-0.71, -0.25] |
| Habitat Breadth | -2.23 | [-2.52, -1.95] |
| Generation Length | 0.43 | [0.20, 0.66] |
| **Categorical Trait** | **Difference** | **Direction** |
| Pelagic Specialists | 19.5% | Greater |
| Diver | -3.2% | Fewer |
| Generalist | -15.3% | Fewer |
| Ground | -5.1% | Fewer |
| Surface | 23.7% | Greater |
| Vertebrates & Scavengers | 33.7% | Greater |
| Invertebrates | -34.1% | Fewer |
| Omnivore | 11.4% | Greater |
| Full Migrants | -0.3% | Fewer |

**Trait redundancy and uniqueness**

To compare the sensitivity of the unique trait combinations (UTC), we remove any species with missing trait values, leaving a total of 281 species for the analysis.

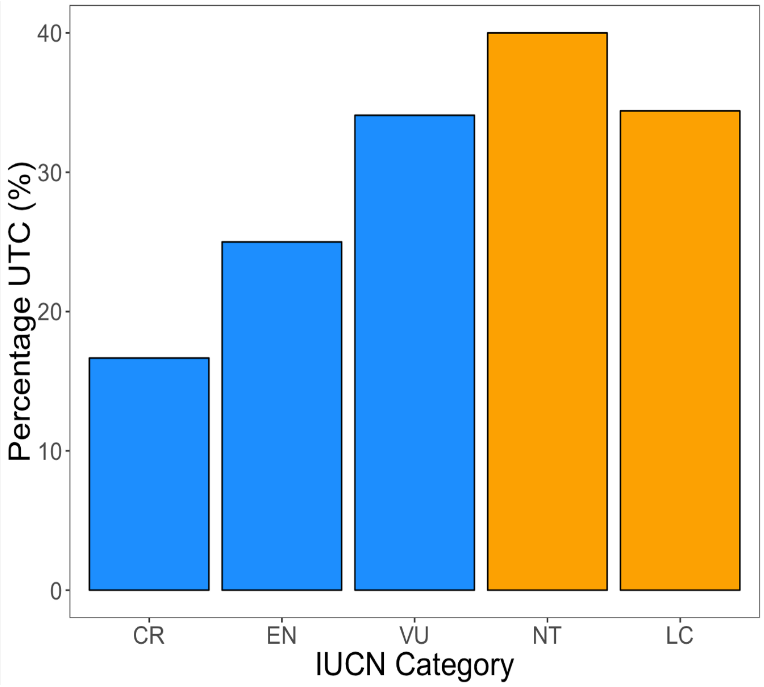


*Figure S2.4 Proportion of seabird species (281 sp.) with non-imputed unique trait combinations for each IUCN category. Orange represents non-threatened categories and blue represents globally threatened categories.*

### SIMPER

*Table S2.3 SIMPER summary of top five non-imputed traits contributing to the Bray Curtis dissimilarity between threats. The proportion of species per trait is indicated as greater (+), or smaller (-) between each threat category. 'S' indicates 'small' and 'M' indicates 'medium'.*

| Threat Contrast | Trait | Contribution (%) | Cumulative (%) | Direct | Habitat | No Threat |
|---|---|---|---|---|---|---|
| *Direct vs. Habitat* | Non-pelagic Specialism | 7.2 | 7.2 | - | + | |
| | Pelagic Specialism | 7.2 | 14.5 | + | - | |
| | Generation Length (S) | 7.1 | 21.6 | - | + | |
| | Clutch Size (S) | 6.7 | 28.3 | + | - | |
| | Generation Length (M) | 6.1 | 34.4 | + | - | |
| *Direct vs. No Threats* | Omnivore Diet | 12.8 | 12.8 | + | | - |
| | Vertebrate and Scavenger | 9.6 | 22.4 | + | | - |
| | Pelagic Specialism | 7.5 | 30.0 | + | | - |
| | Non-pelagic Specialism | 7.5 | 37.5 | - | | + |
| | Habitat Breadth (S) | 6.4 | 43.9 | + | | - |
| *Habitat vs. No Threats* | Omnivore Diet | 11.0 | 11.0 | | + | - |
| | Habitat Breadth (S) | 7.3 | 18.3 | | + | - |
| | Habitat Breadth (M) | 7.1 | 25.4 | | - | + |
| | Vertebrate and Scavenger | 6.4 | 31.8 | | + | - |
| | Clutch Size (M) | 6.3 | 38.0 | | - | + |
